# Exploring the potential of microbial inoculant to enhance common bean (*Phaseolus vulgaris* L.) yield via increased root nodulation and soil macro-nutrients

**DOI:** 10.1101/2025.04.17.649356

**Authors:** Sien Tigah Fortu, Aaron Suh Tening, Solange Takwi Ndzeshala, Christopher Ngosong

## Abstract

Poor soil fertility constraint for common bean (*Phaseolus vulgaris* L.) production is often resolved using chemical fertilizers, but microbial inoculant can be used as sustainable option. Microbial inoculant formulation was tested on common bean in a randomized complete block design field experiment with eight treatments and four replicates. Treatments include untreated control, chemical NPK (nitrogen, phosphorus and potassium), poultry manure (PM), Microbes, NPK+PM, NPK+Microbes, PM+Microbes, and NPK+PM+Microbes. Microbes had higher fertilizer replacement value relative to poultry manure (41.6%) and chemical NPK (26.7%), while poultry manure had 64.3% higher fertilizer replacement value relative to NPK. Bean yield was significantly (*P*<0.001) higher in NPK+PM+Microbes (3.46 tons ha^-1^) and microbial inoculant (1.71 tons ha^-1^) than untreated control (1.42 tons ha^-1^). Higher 1000-grain weight (*P*<0.05) occurred in NPK+PM+Microbes (494 g) and microbes (457 g) than NPK (448 g), PM (436 g) and untreated control (421 g). Effective root nodules correlated positively (*P*<0.05) with bean yield (*r* = 0.77) and 1000-grain weight (*r* = 0.9). Application of NPK+PM+Microbes had higher (9) number of nodules and effectiveness than untreated control (2–3). Treatments increased soil pH significantly (*P*<0.05) in NPK+PM+Microbes and Microbes+PM (6.5) than untreated control (5.5) and NPK (5.0). Soil nitrogen and available phosphorus increased significantly (*P*<0.05) in NPK+PM+Microbes than untreated control, while higher potassium occurred in NPK than untreated control. These findings highlight the potential to explore the ability of bio-inoculant to boost root nodulation, enhance soil macro-nutrients and modulate soil pH buffer in bean production systems.

## 1. Introduction

Ensuring global food and nutrition security requires production of nutrient rich foods such as common bean (*Phaseolus vulgaris* L.), which is a vital source of dietary protein [1,2]. Common bean is widely cultivated for subsistence and commercial purposes, and its production serves a dual role as soil fertility enhancer and food source to over 300 million people [3–5]. Global bean production was 31.8 million tons in 2019, with only 456,000 tons produced in Cameroon at an average yield of 1.23 tons ha^-1^ compared to global average yield of 0.77 tons ha^-1^ and 0.82 tons ha^-1^ in Africa but far below average yield of Mali 10 tons ha^-1^, highest global bean producer per hectare [6]. Low crop yields are partly due to macro-nutrient deficiency (nitrogen–N, phosphorus–P and potassium–K), coupled with pest/disease constraints [7,8].

Various management strategies are being employed to grapple with biotic and abiotic challenges in crop production [3,9,10]. Amongst them, chemical and organic inputs are common, but the indiscriminate use of agro-chemicals threatens the sustainability of agricultural systems due to soil degradation, nutrient imbalance, salinization and acidification, coupled with pest resistance [11,12]. Alternatively, beneficial microbes are being encouraged as sustainable options to improve crop performance including yields of common bean [13], soybean [14] and maize [15], control banana mealybug pest [16], and improve soil physico-chemical properties [17]. Overall, crop yield is largely influenced by soil physico-chemical and biological properties as influenced by farm management practices. Management practices can modulate soil biotic interactions with negative effects on soil biota and nutients that can jeopardize soil sustainability [18–20].

Current calls to harness benefits of the plant microbiome in agriculture is supported by recent studies demonstrating field applications [16,17,21,22]. Microbes can mediate important soil functions such as N fixation, P or K solubilisation and mineralisation [23–25], phytohormones and siderophores production [26,27]. They promote urease activity in soil that supplies N to plants by hydrolysing nitrogenous compounds and boost nitrogenase enzyme complex that fixes atmospheric [17,28]. Plants and microorganisms release phosphatase enzymes that facilitate mobilization of minerally or organically bound phosphates into bioavailable forms and enhance P cycling [29,30]. Microbes also produce siderophore that facilitates solubilization of iron from rhizosphere minerals or organic compounds by binding and reducing Fe^3+^ to Fe^2+^ [27]. Hence, microbial inoculants play vital roles in plant-soil relations that is often ignored but it contributes significantly to crop performances.

Nonetheless, despite positive effects of beneficial microbes, their availability in local markets and adoption is relatively low. Hence, this study aims to evaluate the potential of locally produced microbial inoculant consortium to effectively improve common bean grain yield by enhancing root nodulation and soil macro-nutrient contents. It was hypothesized that the microbial inoculant will increase common bean grain yield via enhanced root nodulation, modulation of soil pH, and enhancement of soil macro-nutrient contents as compared to the untreated chemical or organic amendments and the untreated control.

## 2. Materials and Methods

### 2.1. Experimental site

This experiment was conducted at the Teaching and Research Farm of the Faculty of Agriculture and Veterinary Medicine, University of Buea, Cameroon, between May and August 2021. The site is located at the foot of Mount Cameroon (4100 m) in the South West Region, at about 1000 m above sea level, with coordinates of latitude 04^0^ 8’ 55.1’’ N and longitude 09^0^16’ 53’’ E. Buea has a mono-modal rainfall pattern with rainy season from March to October and 2,085 mm annual average on the leeward side of the mountain and 9,086 mm on the windward side. The dry season runs from November to February, with mean monthly temperatures ranging from 19–30 °C and 28 °C annual mean, with decreasing soil temperature from 25 °C to 15 °C at 10 cm depth as elevation increases from 200–2200 m above sea level (Fraser *et al*., 1998; Proctor *et al*., 2007) [31,32]. The average relative humidity is 85–90%, with annual sunshine ranging between 900–1200 hours, with the baseline soil dominated by sand (68%), silt (18%), and clay (14%), with pH (6.60).

### 2.2. Experimental setup

The field experiment was conducted from May to August 2021 on land that was previously cultivated with cereals (e.g., maize), grain legumes (e.g., cowpea and groundnuts), and vegetables (e.g., huckleberry and amaranth). The land was cleared manually using a cutlass and demarcated into 32 plots (4 m × 4 m). The experiment was set up as randomised complete block design (RCBD) with 8 treatments and 4 replicates. Treatments include an untreated control, chemical NPK, poultry manure, microbes, NPK+poultry manure, NPK+microbes, poultry manure+microbes, and NPK+poultry manure+microbes. The experimental units were separated by 1 m buffer and a 2-m buffer surrounded the experimental site. Soil baseline anaylsis of the experimental site revealed 6.6 pH, 0.25 % N, 8.02 mg/kg P, and 1.62 cmol/kg K (**Table 1**).

**Table 1.** Chemical composition of poultry manure and baseline experimental field soil.

| Treatments | pH(H<sub>2</sub>O) | N<br>(%) | Av. Phosphorus<br>(mg/kg) | Potassium | Calcium | Magnesium |
| --- | --- | --- | --- | --- | --- | --- |
|  |  |  |  | —————cmol/kg————— |  |  |
| Soil | 6.60 ± 0.18b | 0.25 ± 0.02b | 8.02 ± 0.23a | 1.62 ± 0.21b | 2.96 ± 0.30b | 1.24 ± 0.14a |
| Manure | 7.8 ± 0.01a | 3.84 ± 0.01a | 8.35 ± 0.11a | 3.18 ± 0.01a | 5.2 ± 0.01a | 1.37 ± 0.01a |

Values within a column with different letter(s) are significantly different (Tukey's HSD, $P < 0.05$ ).

### 2.3. Preparation of microbial inoculum

A bio-inoculant consortium of beneficial bacteria (**Table 2**) was formulated, comprising symbiotic *Bradyrhizobium japonicum* and *Kosakosania radicincitans,* and non-symbiotic *Arthobacter* spp. (03), *Bacillus* spp. (03), *Lysinibacillus* sp. (01), *Paenibacillus* spp. (02), and *Sinomonas* sp. (01). *B. japonicum* was obtained from the Soil Microbiology Laboratory of the Biotechnology Center of the University of Yaoundé I, Cameroon. *K. radicincitans* isolated from the phyllosphere of winter wheat in Germany, deposited in NCBI as DSM 16656T GenBank: CP018016.1, CP018017.1, CP018018.1, and stored at the Rhizobiology Laboratory (RhizoLab) of the Faculty of Agriculture and Veterinary Medicine of the University of Buea, Cameroon [33,34]. The non-symbiotic bacteria were isolated from the rhizosphere of maize plants in Cameroon [24] and stored at the RhizoLab.

**Table 2.**
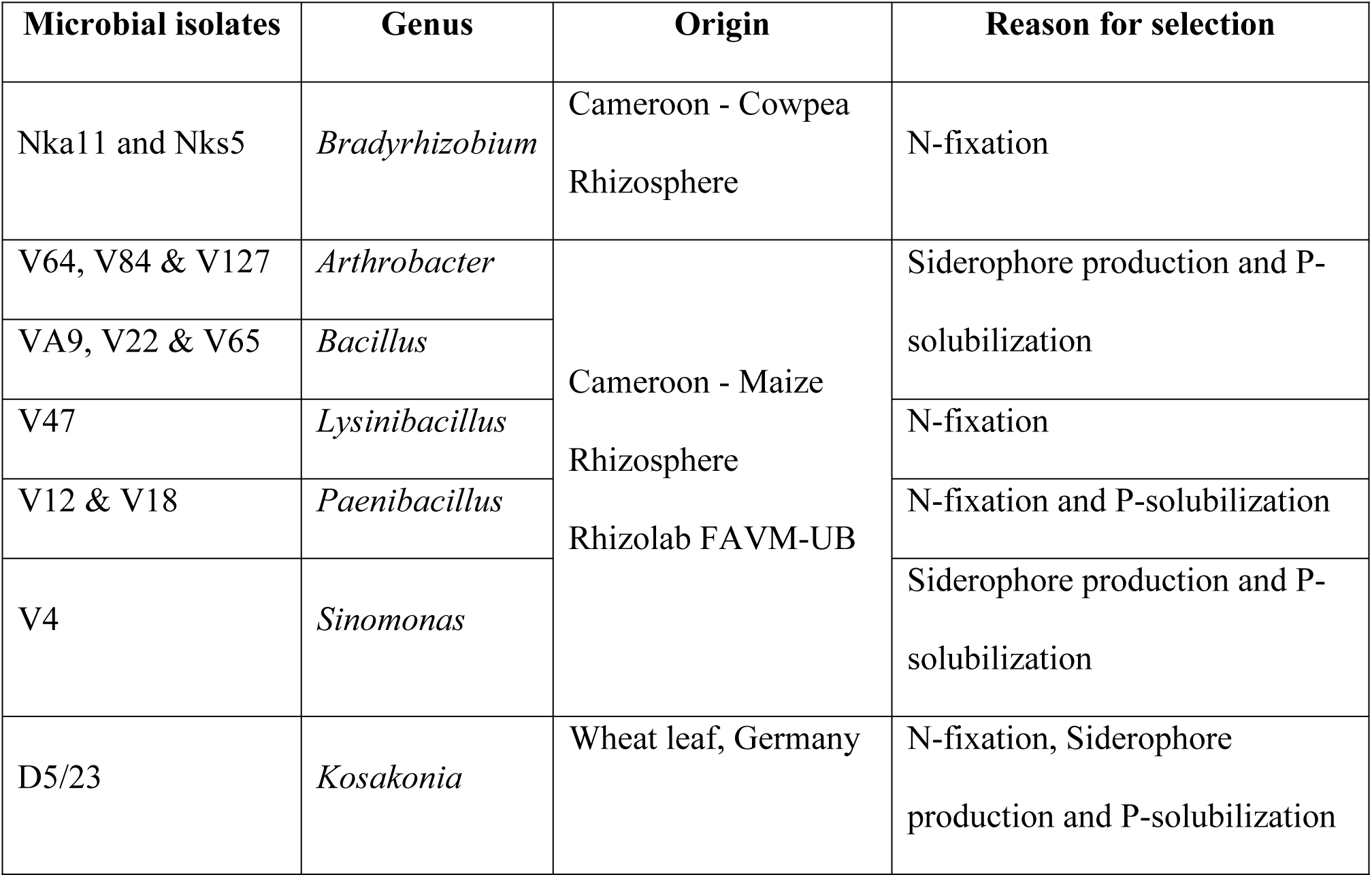
Bacterial strains used in the formulation of microbial inoculant and their potential functions in the inoculant consortium.

A colony of *B. japonicum* was collected from the inoculum stock and transferred into a 500 mL flask containing 100 mL sterilized yeast mannitol broth (YMB), and incubated in a shaker at 28 °C at 200 rpm for 48 hrs. From individual stock culture of the other bacteria, pure colony was collected and transferred into a 500 mL flask containing 100 mL sterilized nutrient broth (Standard nutrient broth I, Carl Roth, Germany), and incubated at 28 °C for 24–48 h. The individually cultured bacteria were assembled into a microbial consortium in a 5 L container. 20.5g of sugar was added to serve as an adjuvant given that *B. japonicum* is not sticky. Common bean seeds were immersed in microbial inoculant consortium (e.g., 1 kg seeds per 100 mL of bio-inoculant) and thoroughly mixed. Seeds were removed from the inoculum and allowed to air-dry for 1 hr before planting [35]. All inoculation procedures were performed under shade to maintain the viability of the microbes.

### 2.4. Soil fertility amendments

The NPK fertilizer (20:10:10 + CaO; ADER^®^ Cameroon) was bought from a local market, while poultry manure (about 2 months old substrate) was collected from a poultry farm in Buea. Granular NPK fertilizer was applied manually by ringing at about 5 cm from plants, as split dose of 5 g NPK per stand at 2 and 4 weeks after sowing [10]. Poultry manure was applied two weeks before planting at 20 tons ha^-1^ [36], giving about 768 kg N, 0.167 kg P and 24.868 kg K in relation to baseline poultry manure anaylsis that had 3.84 % N, 8.35 mg/kg P, and 3.18 cmol/kg K (**Table 1**).

### 2.5. Crop cultivation and management

Untreated bean seeds (ECA PAN 021) were purchased from a local market and sown manually at 35 cm × 35 cm spacing. Two seeds were sown per stand at about 3–4 cm depth and thinned to 1 plant per stand after germination, giving 82,000 plants per hectare. Two weeks before planting, the field was sprayed with pre-emergence systemic herbicide (METROPOLE^®^ - Active ingredient Metribuzin) at 15 mL per 16 L water. After planting, plots were monitored regularly for weed emergence and weeded manually.

### 2.6. Data collection

#### 2.6.1. Soil chemical properties

Pre-planting soil and manure and post-planting (at physiological maturity) soil samples were collected and analyzed for routine parameters (pH, total N, available P, organic carbon, and exchangeable bases) at the Environmental and Analytical Chemistry Laboratory, University of Dschang, Cameroon. Soil samples were collected at 0–15 cm-depth using an auger and stored in polythene bags. Soil and manure samples were air-dried on plastic trays at room temperature, ground, and passed through a 2-mm sieve. Soil pH was determined at 1:2.5 soil/water suspensions using a digital pH meter. Soil available P was determined by the Bray II method [37], organic carbon by the Walkley and Black wet digestion method [38], and total N by the Kjeldahl digestion method [39]. Soil exchangeable bases were extracted with 1 N ammonium acetate (NH_4_OAc) solution at pH 7. After extraction, calcium and magnesium were determined by complexometric titration, while sodium and K were determined using flame photometer [40]. Particle size distribution was determined by the pipette method and textural class assigned according to the USDA textural triangle [37].

#### 2.6.2. Root nodulation

Root nodulation was assessed 65 days after sowing (late flowering) where, ten randomly selected plants per treatment plot were uprooted and placed in a water-filled basin, roots comprising nodules were gently washed with tap water and filtered through a 250-mm sieve. Total number of nodules was counted and all the washed nodules were dissected using a blade to determine the number of effective nodules based on the pigmentation. Nodules with pink or red colouration were recorded as effective in N-fixation due to leghemoglobin oxygen carrier essential for nitrogen fixation, and white nodules were considered as non-effective [41].

#### 2.6.3. Common bean growth and yield parameters

Ten randomly selected plants from the middle rows in each plot were tagged for growth and yield data collection. Plant growth components (plant height, stem girth, leaf area, number of branches, and number of leaves) were collected beginning two weeks after germination and then every week for 6 weeks. Stem girth expressed in centimetres (cm) was measured using a graduated tape at the plant base above the soil surface. Bean yield components number of flowers, number of pods, length of pods, number of seeds per pod were collected before harvest and fresh and dry weight of pods and seeds collected at harvest. Grains were harvested and oven-dried at a constant temperature of 60 °C for three days [42], 1000-grain weight (g) was calculated by taking the weight of randomly selected 1000 grains from each treatment replicate and weighing them, and grain yield was calculated using the following formula [43]:

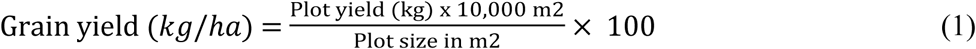

Additionally, the fertilizer replacement value (RV) was calculated as a useful parameter to evaluate the effectiveness of alternative amendment, e.g., comparing microbial inoculant with chemical or organic fertilizers and their combinations. It was calculated based on grain yield using the formula below [44,45]:

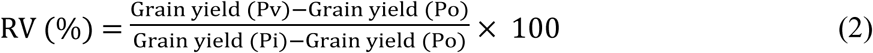

Where Pv refers to microbial inoculant, Po is untreated control, and Pi is chemical or organic fertilizers and the combinations.

### 2.7. Data analysis

All analyses were done using SPSS (Ver. 23) statistical software package, while Microsoft Excel (2019) was used to process graphs and tables. Kolmogorov-Smirnov test was done to test for normality of data. Data for soil physico-chemical properties, root nodulation, growth, and yield parameters were subjected to one-way analysis of variance (ANOVA) to test the effects of treatments. Significant means were separated using Tukey’s Honesty Significant Difference Test (Tukey’s HSD, *P*<0.05). Pearson Correlation (*P*<0.05) was performed to determine the degree of association between variables.

## 3. Results

### 3.1. Fertilizer value of microbial inoculants and bean yield

All parameters analysed were higher in poultry manure over baseline soil (**Table 1**). Poultry manure had higher NPK (3.84 N, 8.35 P and 3.18 K) as compared to the baseline soil (0.25 N, 8.02 P and 1.62 K). Microbial inoculants had higher fertilizer replacement value relative to poultry manure (41.6%) and chemical NPK (26.7%), while poultry manure had 64.3% higher fertilizer replacement value relative to NPK **(Fig. 1**). Common bean yield ranged between 1.42 and 3.46 tons ha^-1^ and increased significantly in all treatments as compared to the untreated control (*P*<0.001, **Fig. 2**). The highest common bean yield was recorded in the integrated application of chemical, organic and microbe (3.46 tons ha^-1^) that differed significantly (*P*<0.001) from all other treatments, with the lowest in untreated control (1.42 tons ha^-1^). Sole application of microbial inoculant increased bean grain yield by 21% as compared to the untreated control, with integrated treatments performing at the rate of 47– 135% better than the untreated control. The weight of 1000 bean grains ranged from 421– 494 g and increased significantly (*P*<0.001, **Fig. 3**) in all treatments compared to the untreated control (421 g), with the highest in the integrated application of chemical, organic and microbe (494 g). Bean yield (*r* = 0.77) and 1000-grain weight (*r* = 0.90) correlated significantly (*P*<0.05) with the number of effective nodules. Integrated application of chemical, organic and microbe resulted in significantly (*P*<0.05, **Table 3**) higher plant height (41.7 cm), as compared to the other treatments with the lowest in untreated control (29.1 cm). The number of leaves followed a similar trend (*P*<0.05, **Table 3**), with the highest in integrated application of chemical, organic and microbe (15) and the lowest in untreated control (11).

**Fig 1.**
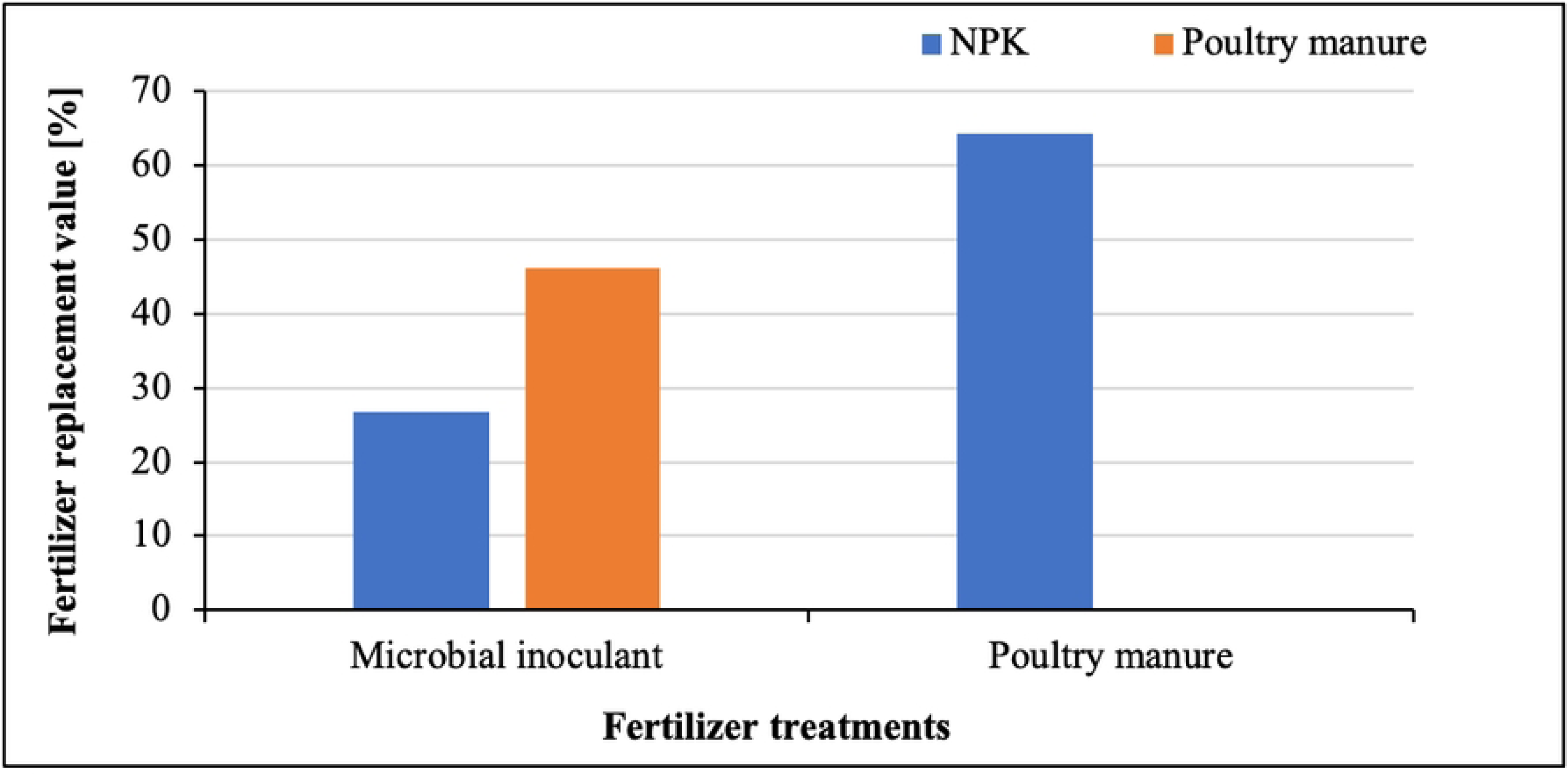
Fertilizer replacement value of microbial inoculant treatment relative to NPK and poultry manure fertilizers, and poultry manure relative to NPK fertilizer.

**Fig 2.**
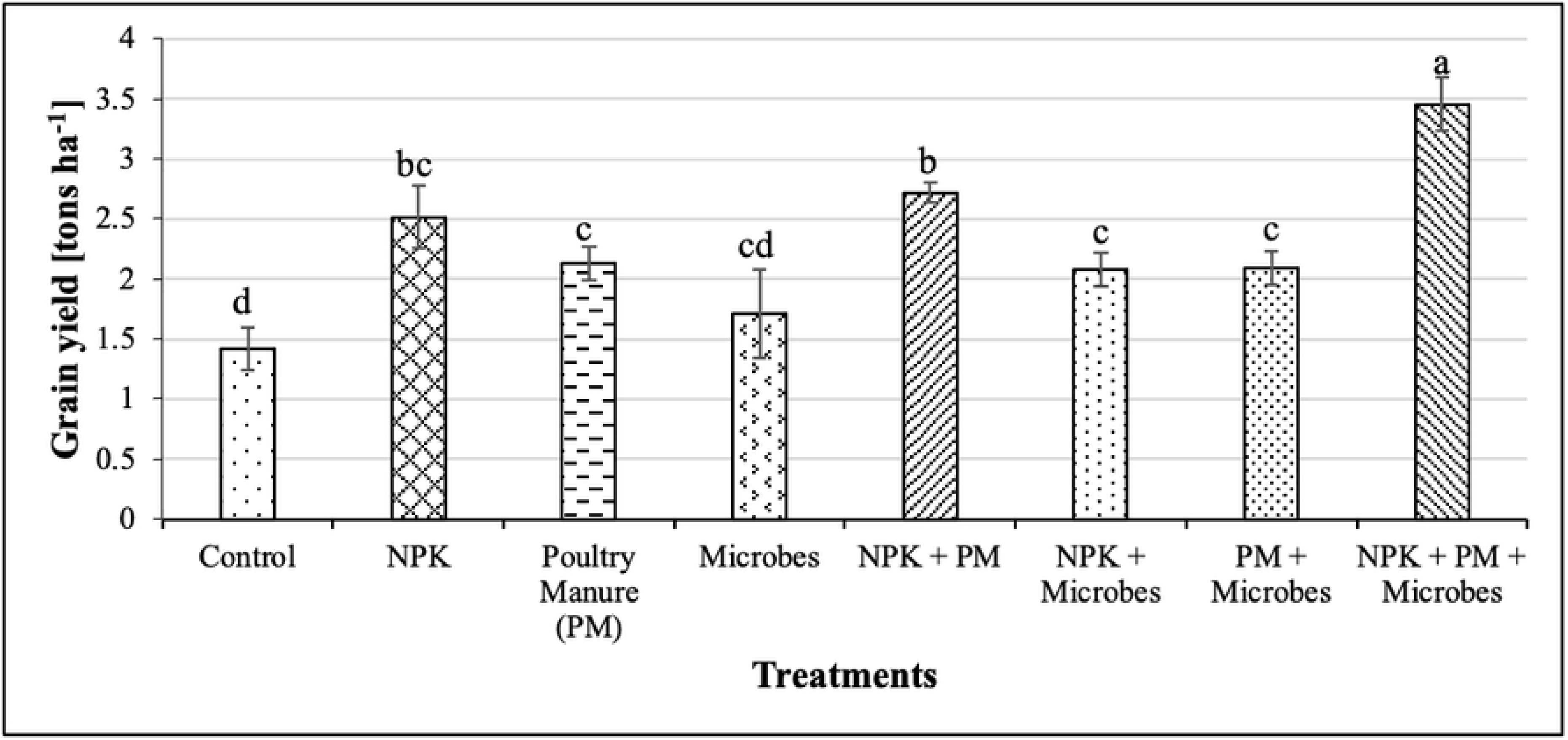
Impact of treatments on common bean grain yield (tons ha^-1^, Mean ± SD). Values with different letters are significantly different (Tukey’s HSD, *P*<0.05).

**Fig 3.**
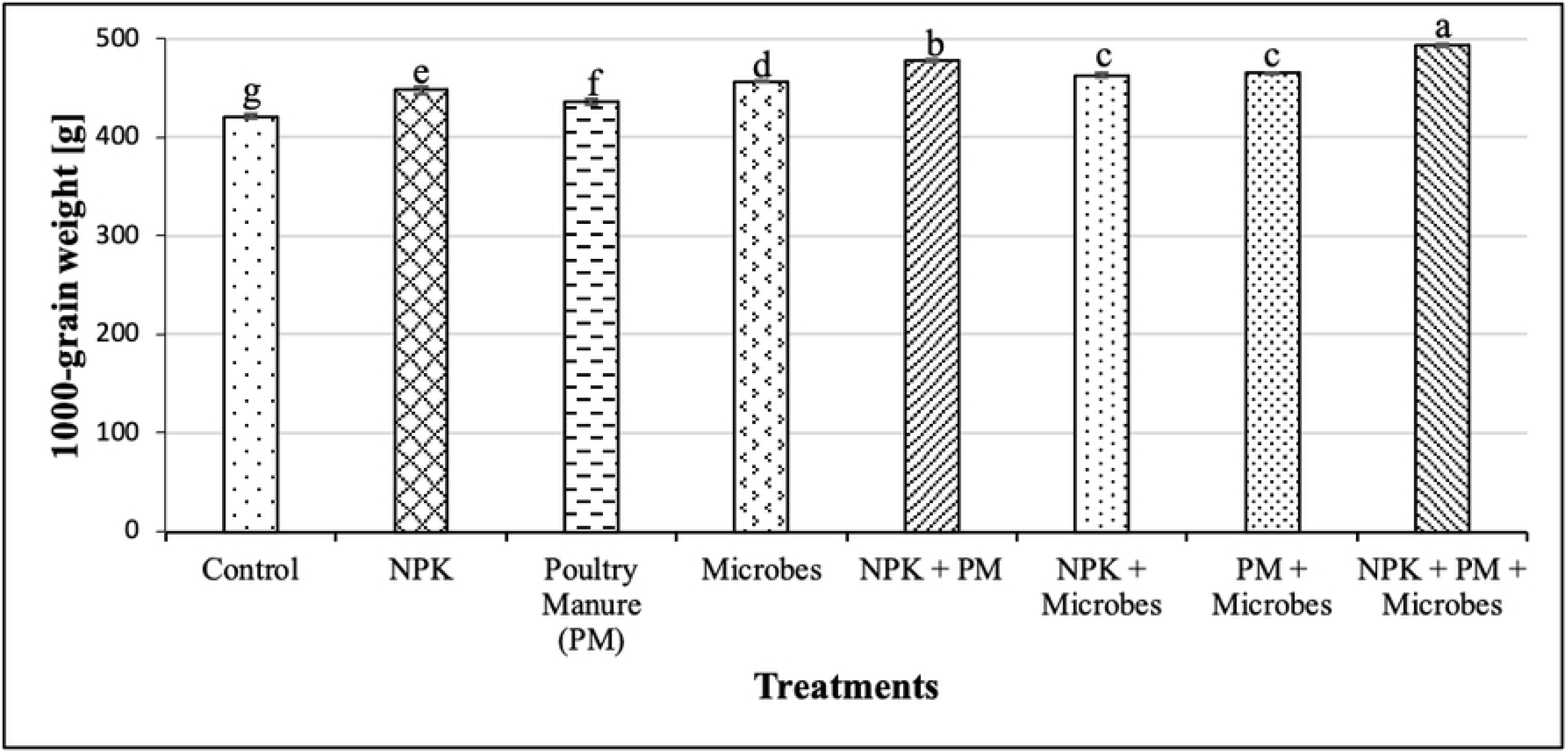
Impact of treatments on 1000-grain weight of common bean (Mean ± SD). Values with different letters are significantly different (Tukey’s HSD, *P*<0.05).

**Table 3.**
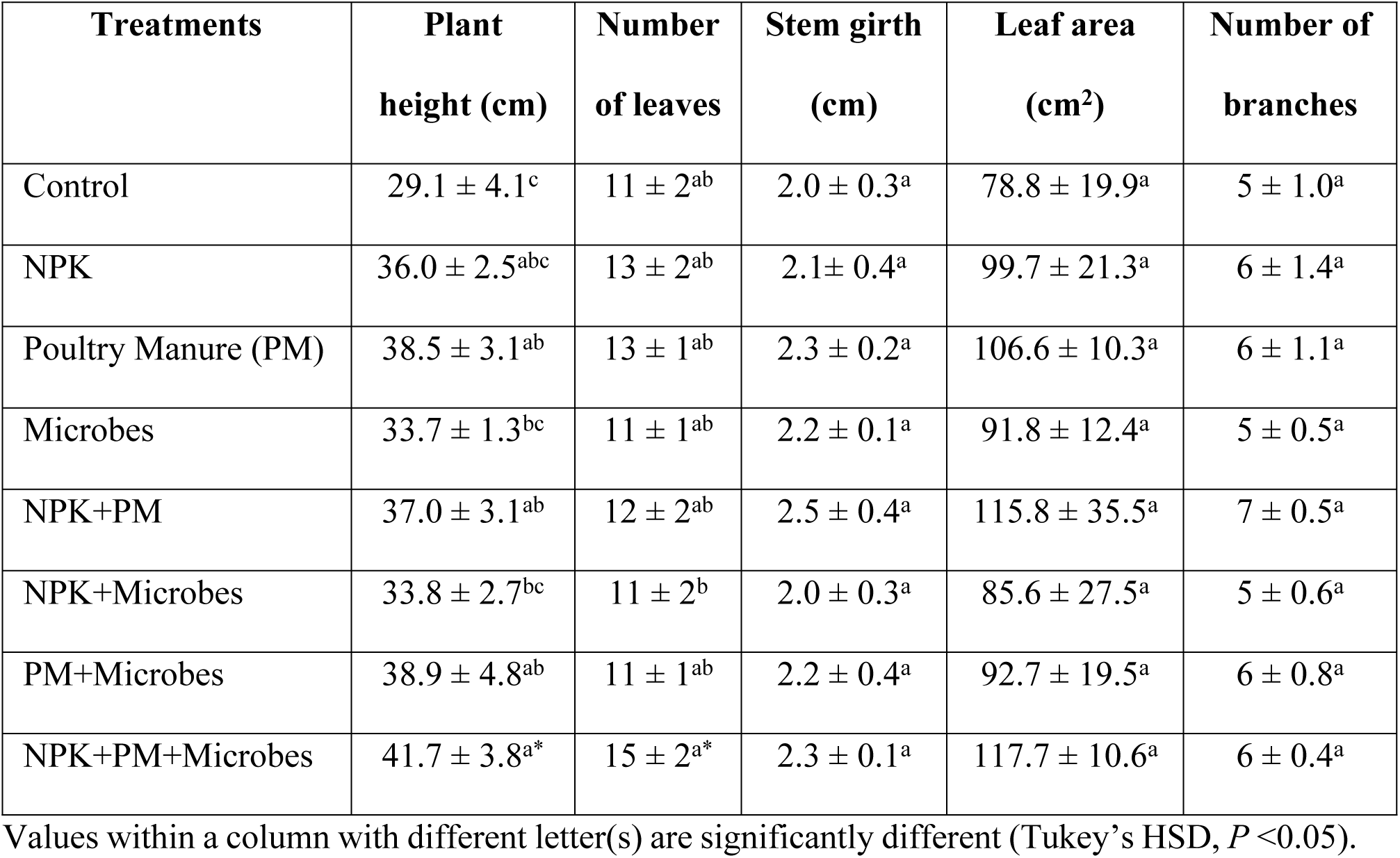
Impact of treatments (Mean ± SD) on plant growth parameters.

### 3.2. Soil chemical properties

Treatments significantly (*P*<0.05) affected the post-planting soil (**Table 4**). Soil pH differed significantly (*P*<0.001) across treatments, with the highest in the integrated application of chemical, organic and microbe (6.53), followed by chemical and microbe (6.43), and the lowest in chemical input (5.0). Total soil N differed significantly (*P*<0.05) across treatments with the highest in the integrated application of chemical, organic and microbe (0.25%) and the lowest in the untreated control (0.13%). Soil available phosphorus also varied significantly (*P*<0.05) across treatments with the highest in the integrated application of chemical, organic and microbe (8.68 mg/kg), and the lowest in the untreated control (6.31 mg/kg). Sole application of NPK recorded the highest (*P*<0.05) potassium (2.8 cmol/kg) compared to the untreated control (1.7 cmol/kg).

**Table 4.**
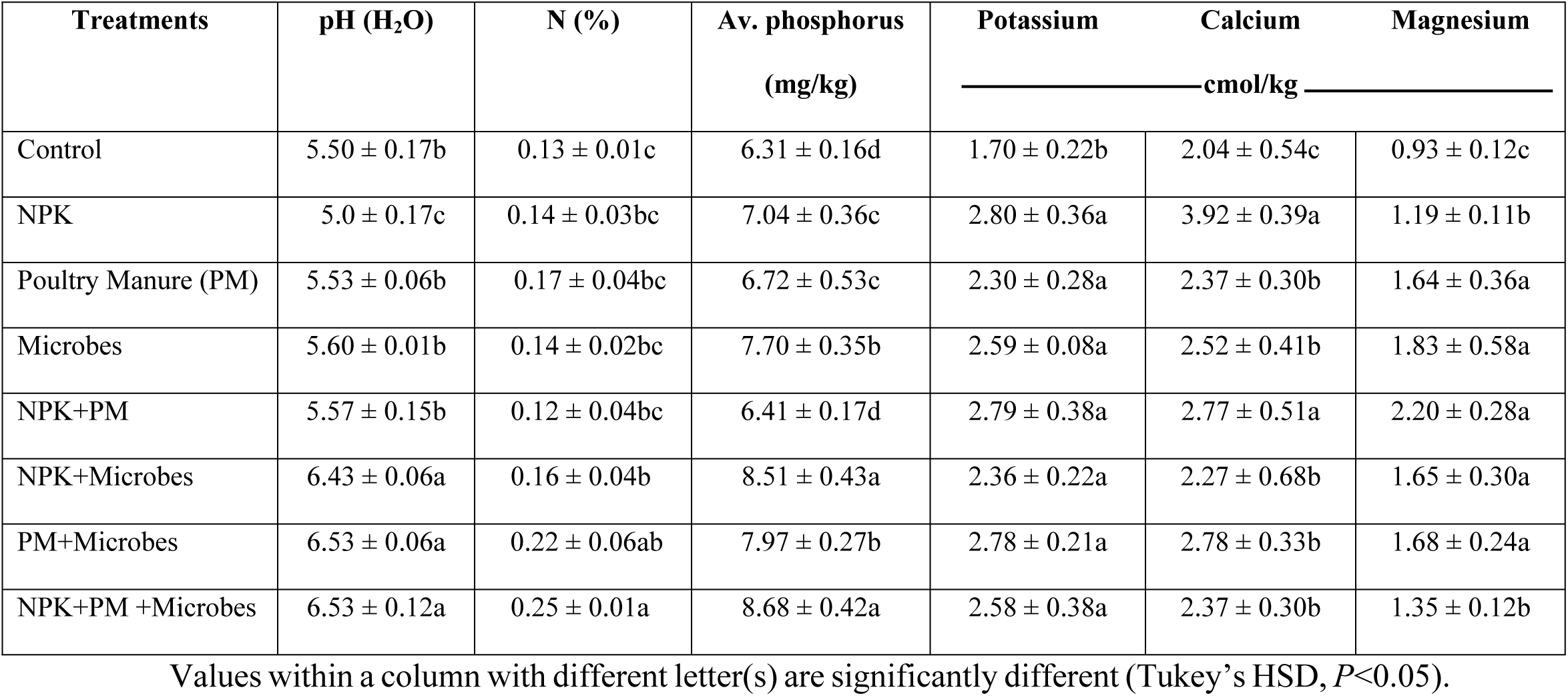
Effect of treatments (Mean ± SD) on soil physico-chemical properties at physiological maturity of common bean.

### 3.3. Root nodulation

Treatments caused significant (*P*<0.05) effects on bean root nodulation parameters (**Fig. 4**), with the highest number of nodules per plant in the integrated application of chemical, organic and microbe (9), which differed significantly from the other treatments and the untreated control (3) with the lowest (*P*<0.001). Similarly, the highest (9) number of effective root nodules per plant was recorded in the integrated application of chemical, organic and microbe treatment and the lowest in untreated control (2). The number of nodules and number of effective nodules also differed significantly (*P*<0.001, **Fig. 5**), and increased in plants inoculated with microbes compared to untreated control.

**Fig 4.**
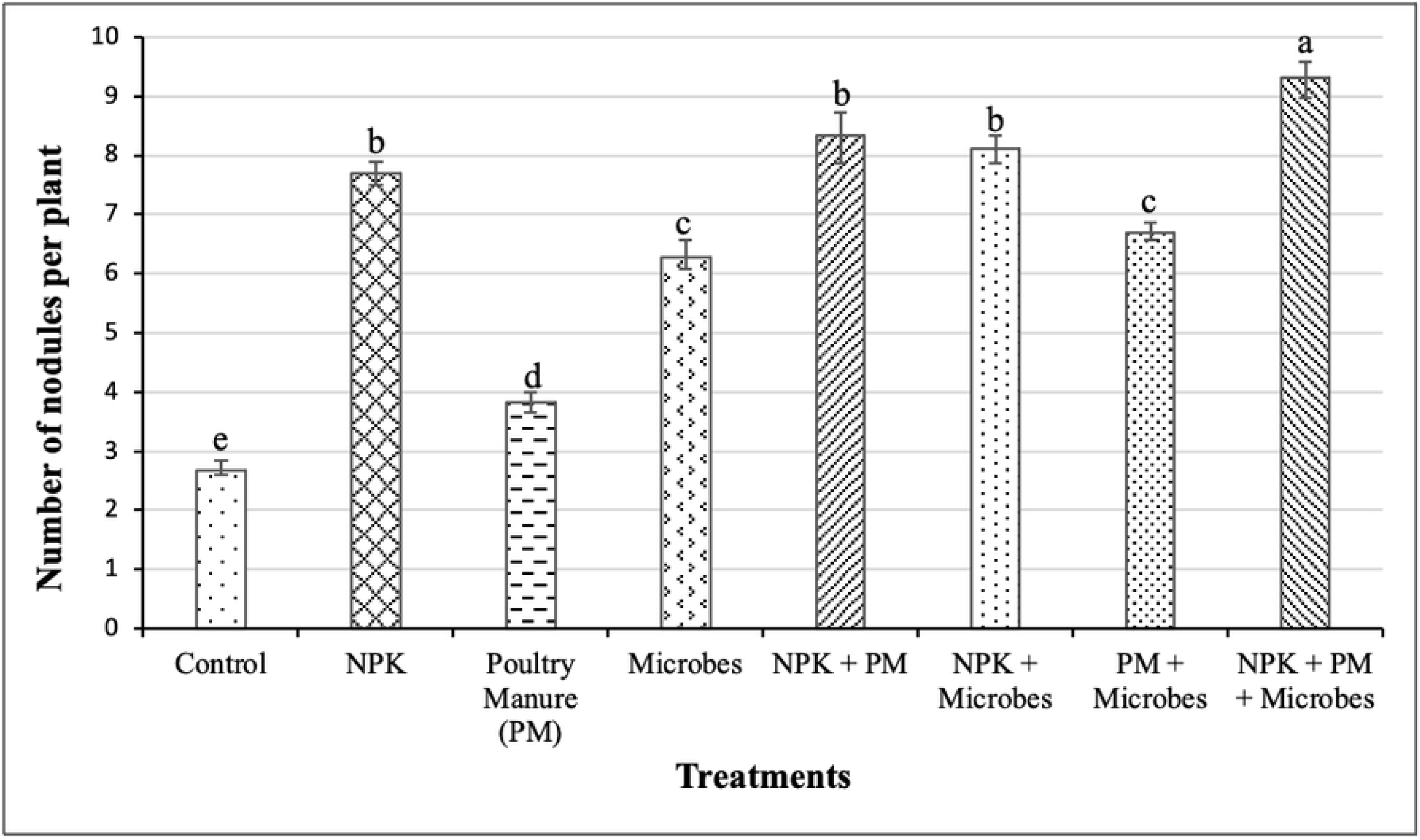
Influence of treatments on number of root nodules (Mean ± SD). Values with different letters are significantly different (Tukey’s HSD, *P*<0.05).

**Fig 5.**
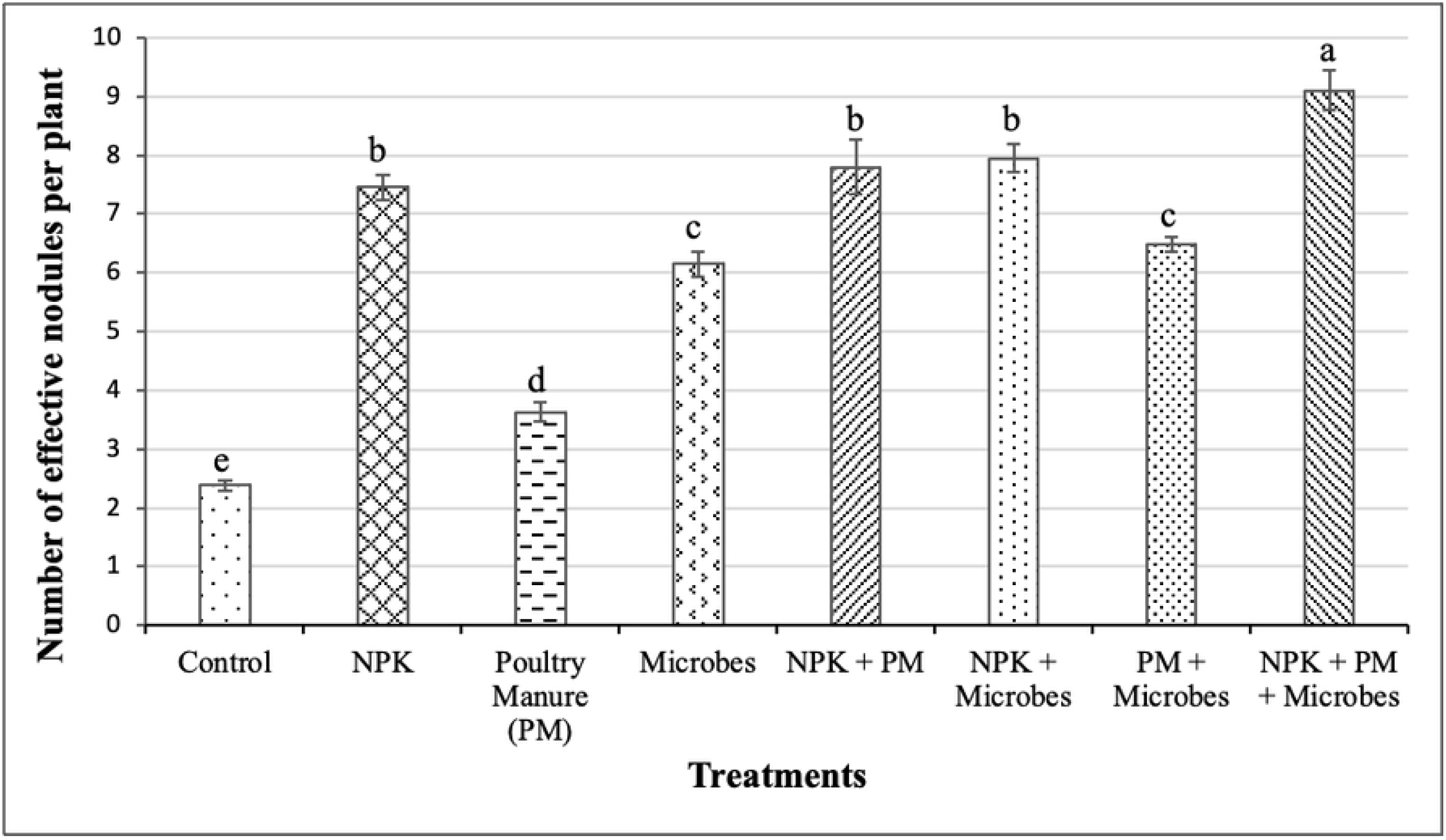
Influence of treatments on number of effective root nodules per plant (Mean ± SD). Values with different letters are significantly different (Tukey’s HSD, *P*<0.05).

## 4. Discussion

### 4.1. Impact of treatments on common bean yield

The higher bean yield for NPK, poultry manure, and microbial inoculant treatments and their integrated application in relation to the untreated control indicates that constant nutrient supply during the crop life cycle boosts productivity [10]. Microbial inoculants can establish themselves in the rhizosphere, facilitating better nutrient acquisition during crop growth, especially in legumes such as common bean, which relies on symbiotic relationships with N-fixing bacteria (*Rhizobium*) to meet their N requirements [46,47]. El-Azeem, [48] reported that co-inoculating *Rhizobium* with *Pseudomonas fluorescens* and siderophore-producing PGPR resulted in enhanced nodulation, plant nutrient uptake and substantial common bean yield component increases like pods per plant and seed weight, compared to sole inoculation. Similarly, Huang *et al*. [49] found that co-inoculating *Bacillus* species with *Rhizobia* led to significant improvements in nodule weight and common bean biomass. Inoculating common bean varieties with *Rhizobium* increases grain yield by enhancing nodulation and improving nitrogen nutrition [50–52]. This aligns with Bojórquez *et al*. [53], who found that co-inoculating *Rhizobium* with PGPB led to improved yield and health of legumes compared to sole *Rhizobium* inoculation. In addition to improving plant nutrient uptake, microbial inoculants (such as *Kosakonia*, *Arthrobacter* and *Bacillus*) can also enhance common bean resilience to environmental stress, thereby contributing to yield stability [13,54]. However, the presence of phosphorus solubilizers and siderophore-producing bacteria such as *Arthrobacter*, *Kosakonia,* and *Sinomonas* in the microbial inoculant consortium likely enhanced phosphorus availability to plants and iron uptake for chlorophyll synthesis and enzyme function [55–57], leading to increased crop performance especially under phosphorus limiting conditions [47,58]. This likely increased photosynthetic efficiency which resulted in improved crop growth and biomass accumulation [57,59]. Increased supplies of nutrients for the integrated application of chemical, organic and microbial inputs resulted in significant increase in nitrogen and phosphorus uptake by plants which eventually influenced common bean yield [60]. The increased symbiotic activity with more N-fixation in plants likely enhanced grain yield [24,34,61] as demonstrated by the significant positive correlation between nodule effectiveness and 1000-grain weight. The poor bean performance in the control treatment emphasizes the need for fertilizer inputs to boost soil fertility and productivity in the study area. In sum, the findings of this study are consistent with the hypothesis that inoculation with beneficial microbes will increase the yield of common beans via enhanced root nodulation and soil fertility status.

### 4.2. Effect of treatments on the soil chemical properties

The enhanced soil fertility with treatment application highlights the role of organic and inorganic fertilizer amendments in improving soil fertility [62,63], and emphasizes the challenges of low soil fertility in cropping systems. The ability of treatments to buffer soil pH, with low values in NPK highlights the ability to cause soil acidification [64–66]. Under acidic pH, crop yield might be reduced as N use efficiency decreases, while excess reactive nitrogen could be lost, thereby, generating H^+^ ions [66,67]. The poultry manure used in this study had a high pH of 7.8 and high Ca^2+^(5.2 %) and Mg^2+^(1.37 %) contents which could have direct and rapid liming effect, proton exchange between soil and organic manure, which often contains some phenolic and humic materials [68,69]. The impacts of poultry manure and microbial applications in modulating the soil pH are consistent with Malik *et al*. [70] and Ogundijo *et al*. [68], who reported significant increase in soil pH with application of organic amendments. PGPBs improve plant nutrient uptake and tolerance to stress, leading to improved yields, and their integration with organic inputs like poultry manure can optimize plant nutrient uptake, boosting productivity and promoting sustainability as seen with the 41.6% fertilizer replacement value of microbial inoculants relative to poultry manure [70,71]. The observed high soil N level for the integrated application of chemical, organic and microbe may be due to high N content of the poultry manure, presence of N fixing bacteria (*Rhizobium*, *Kosakonia, Lysinibacillis* and *Paenibacillus*) in the microbial inoculant consortium and residual N from the chemical fertilizer input [50,68]. High P and K levels also observed in the integrated application of chemical, organic and microbe may be due to high K in poultry manure or *Bacillus* and *Paenibacillus* mediated K uptake by plants [72], residual K from chemical fertilizer input, and solubilizion by *Arthrobacter*, *Bacillus*, *Kosakonia, Paenibacillus* and *Sinomonas* [50,73]. *Arthrobacter*, *Kosakonia,* and *Sinomonas* can enhance plant nutrient solubilization and uptake, improving the overall soil nutrient profile [48,58]. This is important in low-fertility soils where common beans and other legumes are grown. This interaction of PGPB in the soil not only boosts nitrogen levels but also enhances the availability of other essential nutrients such as P and Iron, contributing to better plant health and productivity [74,75]. These findings support the hypothesis of this study that beneficial microbes will modulate soil pH and enhance soil fertility of common bean fields.

### 4.3. Influence of treatments on root nodulation

The observed increase in number of root nodules and their effectiveness in treatments inoculated with microbes is likely due to the influence of *Rhizobium* on N-fixation, which is reflected in the soil N-content that likely resulted in greater photosynthate production for higher grain yield [50,76]. The integrated application of chemical, organic and microbes likely contributed to improved symbiotic N reflected in the high number of effective nodules considering that common bean is a poor N-fixer due to delayed nodule formation with insufficient nodule mass, and ineffective nodules [40,77]. Samago *et al*. [50] and Milcheski *et al*. [78] demonstrated that inoculation with specific rhizobial strains significantly increased nitrogen availability, enhanced nodulation and overall biomass production, leading to higher grain yields. Moreover, common bean has high P-demand for optimum nodulation and growth, which boosts atmospheric N fixation [50,79]. In addition, P initiates nodule formation, increases nodule formation and functions [60,80]. P-solubilizing bacteria in the bio-inoculant consortium such as *Arthrobacter*, *Bacillus*, *Kosakonia, Paenibacillus* and *Sinomonas* likely increased the P-content of soil and enabled optimum soil N-P balance that fostered root nodulation and effectiveness [24,34,61]. The soil also plays a critical role in the effectiveness of microbial inoculants, as the presence of native rhizobia in the soil can enhance their symbiotic relationship with common beans, as demonstrated by Kawaka *et al*. [81], that high populations of native rhizobia in certain soils contributed to effective root nodulation and N fixation. Additionally, P application in combination with rhizobium inoculation increased grain yield, as P availability is essential for N fixation [50,82]. The increase in number of nodules and nodule effectiveness could be attributed to improved nutrient availability such as P and Ca, which might have been supplied by chemical and organic inputs [40,83]. Hence, the decrease in number of nodules under poultry manure compared to chemical fertilizer is probably due to constant nutrient supply by poultry manure, which reduced plant dependence on N-fixation [10,84]. These results support the hypothesis that microbial inoculants will enhance root nodulation and nitrogen fixation, thereby increasing common bean grain yield.

## 5. Conclusion

The locally produced microbial inoculant consortium significantly increased the yield of common bean as compared to the untreated control, while the integrated treatments resulted in higher yields. Thereby, highlighting the importance of microbial inoculants and integrated soil fertility management strategies in increasing common bean yield. Additionally, microbial inoculant consortium and all other treatments increased the number of common bean root nodules and their effectiveness as compared to the control. The observed positive correlation between effective root nodules and common bean grain yield highlights the role of root nodulation in common bean yield. Overall, the locally produced microbial inoculant consortium demonstrated potential as a sustainable option to improve common bean yield, especially when integrated with chemical and organic fertilizer inputs.

## Acknowledgments

We express appreciation to the Faculty of Agriculture and Veterinary Medicine of the University of Buea, and Ministry of Higher Education of Cameroon. We express gratitude to Dr. Olougou Marie Noele Enyoe of the Rhizobiology Laboratory of the University of Buea for the formulation of microbial inoculant consortium.

## References

[1] FAO. (2021a). The State of Food Security and Nutrition in the World 2021. FAO, Rome. http://www.fao.org/state-of-food-security-nutrition/en/

[2] Wudil AH, Usman M, Rosak-Szyrocka J, Pilař L, Boye M. Reversing Years for Global Food Security: A Review of the Food Security Situation in Sub-Saharan Africa (SSA). International Journal of Environmental Research and Public Health. 2022;19(22): 14836. 10.3390/ijerph192214836

[3] Ssekandi W, Mulumba JW, Colangelo P, Nankya R, Fadda C, Karungi J, et al. The use of common bean (*Phaseolus vulgaris*) traditional varieties and their mixtures with commercial varieties to manage bean fly (*Ophiomyia* spp.) infestations in Uganda. Journal of Pest Science. 2015;89(1): 45–57. 10.1007/s10340-015-0678-7

[4] Nanganoa LT, Njukeng JN, Ngosong C, Atache SKE, Yinda GS, Ebonlo JN, et al. Short-Term Benefits of Grain Legume Fallow Systems on Soil Fertility and Farmers Livelihood in the Humid Forest Zone of Cameroon. International Journal of Sustainable Agricultural Research. 2019;6(4): 213–23. 10.18488/journal.70.2019.64.213.223

[5] Andukwa HA, Ntonifor NN. Farmers’ Knowledge and Perception on Common Beans Production Constraints and their Mitigation Methods in the Humid Rainforest and Highland Savanna of Cameroon. Journal of Experimental Agriculture International. 2021;70–85. 10.9734/jeai/2021/v43i230648

[6] FAO. (2021b). World Food and Agriculture - *Statistical Yearbook* 2021. Rome. 10.4060/cb4477en

[7] Karantin DM, George MT, Akwilini JPT. Identification of drought selection indices of common bean (Phaseolus vulgaris L.) genotypes in the Southern Highlands of Tanzania. African Journal of Agricultural Research. 2019;14(3): 161–7. 10.5897/AJAR2018.13370

[8] Halerimana C, Kyamanywa S, Otim MH. Population Dynamics of Selected Field Pests and Their Effect on Grain Yield and Yield Components of Common Bean in Uganda. [Preprint] 2023. Available from: 10.20944/preprints202309.1026.v1

[9] Raimi A, Adeleke R, Roopnarain A. Soil fertility challenges and Biofertiliser as a viable alternative for increasing smallholder farmer crop productivity in sub-Saharan Africa. Moral MT, editor. Cogent Food & Agriculture. 2017;3(1). 10.1080/23311932.2017.1400933

[10] Ngosong C, Nfor IK, Tanyi CB, Olougou MNE, Nanganoa LT, Tening AS. Effect of poultry manure and inorganic fertilizer on earthworms and soil fertility: Implication on root nodulation and yield of climbing bean (*Phaseolus vulgaris*). Fundamental and Applied Agriculture. 2020;5(1): 88–9888–98. 10.5455/faa.76612

[11] Han J, Shi J, Zeng L, Xu J, Wu L. Effects of nitrogen fertilization on the acidity and salinity of greenhouse soils. Environmental Science and Pollution Research International. 2015;22(4): 2976–86. 10.1007/s11356-014-3542-z

[12] Rahman KMA, Zhang D. Effects of Fertilizer Broadcasting on the Excessive Use of Inorganic Fertilizers and Environmental Sustainability. Sustainability. 2018;10(3): 759. 10.3390/su10030759

[13] Horácio EH, Zucareli C, Gavilanes FZ, Yunes JS, Sanzovo AW dos S, Andrade DS. Co-inoculation of rhizobia, azospirilla and cyanobacteria for increasing common bean production. Semina: Ciências Agrárias. 2020;2015–28. 10.5433/1679-0359.2020v41n5supl1p2015

[14] Ngosong C, Tatah BN, Olougou MNE, Suh C, Nkongho RN, Ngone MA, et al. Inoculating plant growth-promoting bacteria and arbuscular mycorrhiza fungi modulates rhizosphere acid phosphatase and nodulation activities and enhance the productivity of soybean (*Glycine max*). Frontiers in Plant Science. 2022;13. 10.3389/fpls.2022.934339

[15] Ngone MA, Ajoacha DMB, Achiri DT, Tchakounté GVT, Ruppel S, Tening AS, et al. Potential of bio-organic amendment of palm oil mill effluent manure and plant growth-promoting bacteria to enhance the yield and quality of maize grains in Cameroon. Soil Security. 2023;11: 100090–0. 10.1016/j.soisec.2023.100090

[16] Becke HI, Achiri TD, Okolle JN, Ntonifor NN, Ruppel S, Ngosong C. Field efficacy of botanicals and beneficial microbes to control banana mealybug (Pseudococcus elisae Borchsenius) (Hemiptera: Pseudococcidae). Crop Protection. 2023;177: 106549. 10.1016/j.cropro.2023.106549

[17] Olougou MNE, Achiri DT, Ngone MA, Ndzeshala SD, Tchakounté GVT, Tening AS, et al. Bio-inoculant consortia modulated plantain (*Musa paradisiaca* L.) growth, rhizosphere pH, acid phosphatase and urease activity. Soil Advances. 2024;100008–8. 10.1016/j.soilad.2024.100008

[18] Qian X, Gu J, Sun W, Li YD, Fu QX, Wang XJ, et al. Changes in the soil nutrient levels, enzyme activities, microbial community function, and structure during apple orchard maturation. Applied Soil Ecology. 2014;77: 18–25. 10.1016/j.apsoil.2014.01.003

[19] Meena RS, Kumar S, Datta R, Lal R, Vijayakumar V, Brtnicky M, et al. Impact of Agrochemicals on Soil Microbiota and Management: A Review. Land. 2020;9(2): 34. 10.3390/land9020034

[20] Cozim-Melges F, Ripoll-Bosch R, Veen GF, Oggiano P, Bianchi FJJA, van der Putten WH, et al. Farming practices to enhance biodiversity across biomes: a systematic review. npj Biodiversity. 2024;3(1): 1–11. 10.1038/s44185-023-00034-2

[21] Reed L, Glick BR. The recent use of plant-growth-promoting bacteria to promote the growth of agricultural food crops. Agriculture. 2023;13(5): 1089. 10.3390/agriculture13051089

[22] Rios-Ruiz WF, Tuanama-Reátegui C, Huamán-Córdova G, Valdez-Nuñez RA. Co-inoculation of endophytes *Bacillus siamensis* TUR07-02b and *Priestia megaterium* SMBH14-02 promotes growth in rice with low doses of nitrogen fertilizer. Plants. 2023;12(3): 524–4. 10.3390/plants12030524

[23] Abbasi KM, Manzoor M. Bio-solubilization of phosphorus from rock phosphate and other P fertilizers in response to phosphate solubilizing bacteria and poultry manure in a silt loam calcareous soil. Journal of Plant Nutrition and Soil Science. 2018;181(3): 345–56. 10.1002/jpln.201800012

[24] Tchakounté GVT, Berger B, Patz S, Henri F, Ruppel S. Community structure and plant growth-promoting potential of cultivable bacteria isolated from Cameroon soil. Microbiology Research. 2018;214: 47–59. 10.1016/j.micres.2018.05.008

[25] Farhat F, Tariq A, Waseem M, Masood A, Raja S, Ajmal W, et al. Plant Growth Promoting Rhizobacteria (PGPR) Induced Improvements in the Growth, Photosynthesis, Antioxidants, and Nutrient Uptake of Rapeseed (*Brassica napus* L.). Gesunde Pflanzen. 2023;2075–2088. 10.1007/s10343-023-00845-0

[26] Radhakrishnan R, Hashem A, Abd_Allah EF. *Bacillus*: A Biological Tool for Crop Improvement through Bio-Molecular Changes in Adverse Environments. Frontiers in Physiology. 2017;8. 10.3389/fphys.2017.00667

[27] Sultana S, Alam S, Karim MM. Screening of siderophore-producing salt-tolerant rhizobacteria suitable for supporting plant growth in saline soils with iron limitation. Journal of Agriculture and Food Research. 2021;4: 100150. 10.1016/j.jafr.2021.100150

[28] Cordero I, Snell H, Bardgett RD. High throughput method for measuring urease activity in soil. Soil Biology and Biochemistry. 2019;134: 72–7. 10.1016/j.soilbio.2019.03.014

[29] Behera BC, Yadav H, Singh SK, Mishra RR, Sethi BK, Dutta SK, et al. Phosphate solubilization and acid phosphatase activity of Serratia sp. isolated from mangrove soil of Mahanadi river delta, Odisha, India. Journal of Genetic Engineering and Biotechnology. 2017;15(1): 169–78. 10.1016/j.jgeb.2017.01.003

[30] Cabugao KG, Timm CM, Carrell AA, Childs J, Lu TYS, Pelletier DA, et al. Root and Rhizosphere Bacterial Phosphatase Activity Varies with Tree Species and Soil Phosphorus Availability in Puerto Rico Tropical Forest. Frontiers in Plant Science. 2017;8. 10.3389/fpls.2017.01834

[31] Fraser PJ, Hall JB, Healey JR. Climate of the Mount Cameroon Region, long and medium term rainfall, temperature and sunshine data. SAFS, University of Wales Bangor, MCP-LBG. Limbe. 1998;56.

[32] Proctor J, Edwards ID, Payton RW, Nagy L. Zonation of forest vegetation and soils of Mount Cameroon, West Africa. Plant Ecology. 2007;192(2): 251–69. 10.1007/s11258-007-9326-5

[33] Ruppel S, Merbach W. Effects of different nitrogen sources on nitrogen fixation and bacterial growth of Pantoea agglomerans and Azospirillum sp. in bacterial pure culture: An investigation using 15N2 incorporation and acetylene reduction measures. Microbiological Research. 1995;150(4): 409–18. 10.1016/S0944-5013(11)80023-6

[34] Berger B, Patz S, Ruppel S, Dietel K, Faetke S, Junge H, et al. Successful Formulation and Application of Plant Growth-Promoting *Kosakonia radicincitans* in Maize Cultivation. BioMed Research International. 2018;2018: e6439481. 10.1155/2018/6439481

[35] Atieno M, Herrmann L, Okalebo R, Lesueur D. Efficiency of different formulations of *Bradyrhizobium japonicum* and effect of co-inoculation of *Bacillus subtilis* with two different strains of *Bradyrhizobium japonicum*. World Journal of Microbiology and Biotechnology. 2012;28(7): 2541–50. 10.1007/s11274-012-1062-x

[36] Alhrout H, Khalifeh H, Dalaeen H. The impact of organic and inorganic fertilizer on yield and yield components of common bean (*Phaseolus vulgaris*). Advances in Environmental Biology. 2016; 10(9): 8–13.

[37] Van Reeuwijk LP. Procedures for Soil Analysis, 3rd edn, International Soil Reference and Information Centre (ISRIC), Wageningen, 1992.

[38] Kalra YP, Maynard DG. Methods manual for forest soil and plant analysis. Forestry Canada, Northwest Region, Northern Forestry Centre, Edmonton, Alberta. Information Report NOR-X-319E. 1991; 116 p.

[39] Bremner JM, Mulvaney CS. Nitrogen-Total. In: Methods of soil analysis. Part 2. Chemical and microbiological properties. Agronomy Monographs. 1982;595–624. 10.2134/agronmonogr9.2.2ed.c31

[40] Rowell DL. Soil Science: Methods & Applications (1st ed.). Routledge. 1994. 10.4324/9781315844855

[41] Shanka D, Dechassa N, Gebeyehu S, Elias E. Dry matter yield and nodulation of common bean as influenced by phosphorus, lime and compost application at Southern Ethiopia. Open Agriculture. 2018;3(1): 500–9. 10.1515/opag-2018-0055

[42] Doymaz İ. Hot-Air Drying and Rehydration Characteristics of Red Kidney Bean Seeds. Chemical Engineering Communications. 2015;203(5): 599–608. 10.1080/00986445.2015.1056299

[43] FAO and DWFI. Yield gap analysis of field crops – Methods and case studies, by V.O. Sadras, K.G.G. Cassman, P. Grassini, A.J. Hall, W.G.M. Bastiaanssen, A.G. Laborte, A.E. Milne, G. Sileshi, P. Steduto, FAO Water Reports No. 41, Rome, Italy, 2015.

[44] Ashekuzzaman SM, Forrestal P, Richards KG, Daly K, Fenton O. Grassland phosphorus and nitrogen fertiliser replacement value of dairy processing dewatered sludge. Sustainable Production and Consumption. 2021;25: 363–73. 10.1016/j.spc.2020.11.017

[45] Ayeyemi T, Recena R, García-López AM, Delgado A. Circular economy approach to enhance soil fertility based on recovering phosphorus from wastewater. Agronomy. 2023;13(6): 1513. 10.3390/agronomy13061513

[46] Behr JH, Kampouris ID, Babin D, Sommermann L, Francioli D, Kuhl-Nagel T, et al. Beneficial microbial consortium improves winter rye performance by modulating bacterial communities in the rhizosphere and enhancing plant nutrient acquisition. Frontiers in Plant Science. 2023;14. 10.3389/fpls.2023.1232288

[47] Xavier GR, Jesus E da C, Dias A, Coelho MRR, Molina YC, Rumjanek NG. Contribution of biofertilizers to pulse crops: From single-strain inoculants to new technologies based on microbiomes strategies. Plants. 2023;12(4): 954. 10.3390/plants12040954

[48] El-Azeem SAEMA, Effects of co-inoculation of rhizobium and plant growth-promoting rhizobacteria on common bean (*Phaseolus vulgaris*) yield, nodulation, nutrient uptake, and microbial activity under field conditions. Journal of Soil and Water Sciences. 2022;7(1): 13–26. 10.21608/jsws.2022.290563

[49] Huang Z, Cui C, Cao Y, Dai J, Cheng X, Hua S, et al. Tea plant–legume intercropping simultaneously improves soil fertility and tea quality by changing *Bacillus* species composition. Horticulture Research. 2022;9. 10.1093/hr/uhac046

[50] Samago TY, Anniye EW, Dakora FD. Grain yield of common bean (*Phaseolus vulgaris* L.) varieties is markedly increased by rhizobial inoculation and phosphorus application in Ethiopia. Symbiosis. 2017;75(3): 245–55. 10.1007/s13199-017-0529-9

[51] Karavidas I, Ntatsi G, Ntanasi T, Vlachos I, Tampakaki A, Iannetta PPM, et al. Comparative assessment of different crop rotation schemes for organic common bean production. Agronomy. 2020;10(9): 1269. 10.3390/agronomy10091269

[52] Reis DR, Teixeira GC, Teixeira IR, Silva GR, Ribeiro BB. Organomineral fertilization associated with inoculation of *Rhizobium tropici* and co-Inoculation of *Azospirillum brasilense* in common bean. Sustainability. 2023;15(24): 16631–1. 10.3390/su152416631

[53] Armenta-Bojórquez AD, Roblero-Ramírez HR, Camacho-Báez JR, Mundo-Ocampo M, García-Gutiérrez C, Armenta-Medina A. Organic versus synthetic fertilisation of beans (*Phaseolus vulgaris* L.) in Mexico. Experimental Agriculture. 2015;52(1): 154–62. 10.1017/S0014479715000010

[54] Benmrid B, Ghoulam C, Zeroual Y, Kouisni L, Bargaz A. Bioinoculants as a means of increasing crop tolerance to drought and phosphorus deficiency in legume-cereal intercropping systems. Communications Biology. 2023;6(1): 1–15. 10.1038/s42003-023-05399-5

[55] Alori ET, Glick BR, Babalola OO. Microbial phosphorus solubilization and its potential for use in sustainable agriculture. Frontiers in Microbiology. 2017;8. 10.3389/fmicb.2017.00971

[56] Prabhu N, Borkar S, Garg S. Phosphate solubilization by microorganisms. Advances in Biological Science Research. 2019;161–76. 10.1016/B978-0-12-817497-5.00011-2

[57] Cui K, Xu T, Chen J, Yang H, Liu X, Zhuo R, et al. Siderophores, a potential phosphate solubilizer from the endophyte *Streptomyces* sp. CoT10, improved phosphorus mobilization for host plant growth and rhizosphere modulation. Journal of Cleaner Production. 2022;367: 133110–0. 10.1016/j.jclepro.2022.133110

[58] Pattnaik S, Mohapatra B, Gupta A. Plant growth-promoting microbe mediated uptake of essential nutrients (Fe, P, K) for crop stress management: Microbe–Soil–Plant Continuum. Frontiers in Agronomy. 2021;3. 10.3389/fagro.2021.689972

[59] Mortinho ES, Jalal A, Eduardo C, Fernandes GC, Pereira NC, Rosa PA, et al. Co-inoculations with plant growth-promoting bacteria in the common bean to increase efficiency of NPK fertilization. Agronomy. 2022;12(6): 1325–5. 10.3390/agronomy12061325

[60] Zafar M, Abbasi MK, Rahim N, Khaliq A, Shaheen A, Jamil M, et al. Influence of integrated phosphorus supply and plant growth promoting rhizobacteria on growth, nodulation, yield and nutrient uptake in *Phaseolus vulgaris*. African Journal of Biotechnology. 2011;10(74). 10.5897/AJB11.1395

[61] Tchakounté GVT, Berger B, Patz S, Becker M, Turečková V, Ondřej N, et al. The response of maize to inoculation with *Arthrobacter* sp. and *Bacillus* sp. in Phosphorus-Deficient, Salinity-Affected Soil. Microorganisms. 2020;8(7): 1005–5. 10.3390/microorganisms8071005

[62] Mahmood F, Khan I, Ashraf U, Shahzad T, Hussain S, Shahid M, et al. Effects of organic and inorganic manures on maize and their residual impact on soil physicochemical properties. Journal of soil science and plant nutrition. 2017;17(1): 22–32. 10.4067/S0718-95162017005000002

[63] Li X, Su Y, Ahmed T, Ren H, Javed MR, Yao Y, et al. Effects of different organic fertilizers on improving soil from newly reclaimed land to crop soil. Agriculture. 2021;11(6): 560. 10.3390/agriculture11060560

[64] Liu X, Shi H, Bai Z, Liu X, Yang B, Yan D. Assessing soil acidification of croplands in the Poyang Lake Basin of China from 2012 to 2018. Sustainability. 2020;12(8): 3072. 10.3390/su12083072

[65] Liu Y, Zhang M, Li Y, Zhang Y, Huang X, Yang Y, et al. Influence of nitrogen fertilizer application on soil acidification characteristics of tea plantations in Karst areas of Southwest China. Agriculture. 2023;13(4): 849. 10.3390/agriculture13040849

[66] Tkaczyk P, Mocek-Płóciniak A, Skowrońska M, Bednarek W, Kuśmierz S, Zawierucha E. The mineral fertilizer-dependent chemical parameters of soil acidification under field conditions. Sustainability. 2020;12(17): 7165. 10.3390/su12177165

[67] Govindasamy P, Muthusamy SK, Bagavathiannan M, Mowrer J, Jagannadham PT, Maity A, et al. Nitrogen use efficiency—A key to enhance crop productivity under a changing climate. Frontiers in Plant Science. 23;14: 1121073. 10.3389/fpls.2023.1121073

[68] Ogundijo D, Adetunji M, Azeez J, T. Arowolo, Olla N, Adekunle A. Influence of organic and inorganic fertilizers on soil chemical properties and nutrient changes in an Alfisol of South Western Nigeria. International Journal of Plant & Soil Science. 2015;7(6): 329–37. 10.9734/IJPSS/2015/18355

[69] Masocha BL, Dikinya O. The role of poultry litter and its biochar on soil fertility and *Jatropha curcas* L. growth on sandy-loam soil. Applied Sciences. 2022;12(23): 12294. 10.3390/app122312294

[70] Meza C, Valenzuela F, Echeverrı’a-Vega A, Gomez A, Sarkar S, Cabeza RA, et al. Plant-growth-promoting bacteria from rhizosphere of Chilean common bean ecotype (*Phaseolus vulgaris* L.) supporting seed germination and growth against salinity stress. Frontiers in Plant Science. 2022;13: 1–18. 10.3389/fpls.2022.1052263

[71] Raghavan RS, Vidya P, Balakumaran MD, Ramya GK, Nithya K. Plant growth-promoting bacteria: a catalyst for advancing horticulture applications. Biosciences Biotechnology Research Asia. 2024;21(3). 10.13005/bbra/3276

[72] Gundlach J, Herzberg C, Kaever V, Gunka K, Hoffmann T, Weiss MH. Control of potassium homeostasis is an essential function of the second messenger cyclic di-amp in *Bacillus subtillis*. Science Signaling. 2017;10(475): eaal3011. 10.1126/scisignal.aal3011

[73] Dhaliwal SS, Sharma V, Shukla AK, Gupta RK, Verma V, Kaur M, et al. Residual effect of organic and inorganic fertilizers on growth, yield and nutrient uptake in wheat under a basmati rice-wheat cropping system in North-Western India. Agriculture. 2023;13(3): 556. 10.3390/agriculture13030556

[74] Cardoso AM, da Silva CVF, de Pádua VL. Microbial insights into biofortified common bean cultivation. Science. 2024;6(1): 6. 10.3390/sci6010006

[75] Nabi M. Role of microorganisms in plant nutrition and soil health. Sustainable Plant Nutrition. 2023;263–282. 10.1016/B978-0-443-18675-2.00016-X

[76] Shome S, Barman A, Solaiman ZM. Rhizobium and phosphate solubilizing bacteria influence the soil nutrient availability, growth, yield, and quality of soybean. Agriculture. 2022;12(8): 1136. 10.3390/agriculture12081136

[77] Reinprecht Y, Schram L, Marsolais F, Smith TH, Hill B, Pauls KP. Effects of nitrogen application on nitrogen fixation in common bean production. Frontiers in Plant Science. 2020;11: 534817. 10.3389/fpls.2020.01172

[78] Milcheski VF, Senff SE, Orsi N, Botelho GR, Fioreze ACCL. Influence of common bean genotypes and rhizobia interaction for nodulation and nitrogen fixation. Journal of Agroveterinary Sciences. 2022;21(1): 8–15. 10.5965/223811712112022008

[79] Silva DA, Esteves JAD, Messias UA, Teixeir A, Goncalves JGR, Chiorato AF, et al. Efficiency in the use of phosphorus by common bean genotypes. Scientia Agricola. 2014;71(3): 232–239. 10.1590/S0103-90162014000300008

[80] Li H, Wang X, Liang Q, Lyu X, Li S, Gong Z, et al. Regulation of phosphorus supply on nodulation and nitrogen fixation in soybean plants with dual-root systems. Agronomy. 2021;11(11): 2354. 10.3390/agronomy11112354

[81] Kawaka F, Dida MM, Opala PA, Ombori O, Maingi JM, Osoro N, et al. Symbiotic efficiency of native rhizobia nodulating common bean (*Phaseolus vulgaris* L.) in soils of western Kenya. International Scholarly Research Notices. 2014;(1): 258497. 10.1155/2014/258497

[82] Jalal A, Oliveira CE, Bastos AD, Fernandes GC, De Lima BH, Junior EF, et al. Nanozinc and plant growth-promoting bacteria improve biochemical and metabolic attributes of maize in tropical Cerrado. Frontiers in Plant Science. 2023;13: 1046642. 10.3389/fpls.2022.1046642

[83] Endalkachew F, Kibebew K, Asmare M, Bobe B. Yield of faba bean (*Vicia faba* L.) as affected by lime, mineral P, farmyard manure, compost and *Rhizobium* in acid soil of Lay Gayint District, northwestern highlands of Ethiopia. Journal of Agriculture and Food Security. 2018;7: 16. 10.1186/s40066-018-0168-2

[84] Anuruddi HIGK, Ekanayake EMHGS, Kumara RKGK, Kulasooriya SA, Fonseka DLCK. Evaluation of Rhizobial inoculation in comparison to urea fertilizer application of vegetable bean (*Phaseolus vulgaris* L.) Journal of the University of Ruhuna. 2023;11(1): 15–22. 10.4038/jur.v11i1.7987

